# Black Rice Developed Through Interspecific Hybridization (*O. sativa* x *O. rufipogon*): Origin of Black Rice Gene from Indian Wild Rice

**DOI:** 10.1101/2020.12.25.423663

**Authors:** Subhas Chandra Roy, Pankaj Shil

## Abstract

Rice (*Oryza sativa* L.) is a most important staple food grain consumed by more than half of the world’s population. Wild rice (*O. rufipogon* Griff.) is considered as the immediate ancestral progenitor of cultivated rice *O. sativa*, evolved through the process of domestication. Most of the cultivated rice produced grains with white pericarp, but can also produce grains with brown, red and black (or purple rice) pericarp. Red rice pericarp accumulates proanthocyanidin whereas black rice contains anthocyanin, both have antioxidant activity and health benefits. Black pericarp is predicted to be regulated by alleles of three genetic loci- Kala1, Kala3, and Kala4. Recombinational and insertional genetic rearrangement in the promoter region of Kala4 is crucial for the development of black pericarp in rice grain. In the present study, we report first time in the breeding history that aromatic black rice lines were developed through interspecific hybridization and introgression in the genetic background of *O. sativa* **cv.** Badshabhog, Chenga and Ranjit. Badshabhog and Ranjit is white grain rice but Chenga is red rice category. Common Asian wild rice *O. rufipogon* is used as donor parent (red grain) and source of black rice gene. Several possible genetic explanations have come up for the creation of black rice pericarp in the progeny lines. Possible reason may be the rearrangement and insertion of LINE1 in the promoter region of Kala4 allele through recombination mechanism leading to ectopic expression of Kala4 gene for the accumulation of anthocyanin and resulted in black rice formation. Other genes and regulatory factors may be induced and become functional to produce black pericarp. Black pericarp colour appeared in F2 populations in the wide crosses (Badshabhog x *O. rufipogon* and Chenga x *O. rufipogon*) but not in the cross with (Ranjit x *O. rufipogon*). Black pericarp trait inherited in F4 and F5 population with segregation phenotypes.

This is a first report in the history of rice genetics and pre-breeding research, that black rice has been created through wide crossing and introgression by combining wild rice *O. rufipogon* in the genetic background of *O. sativa*. Present experimental evidence provides a new model of black rice origin. Thus, black rice (indica type) of Indian subcontinent originated independently through natural out crossing and artificial selection in the course of domestication.

## Introduction

Rice (*Oryza sativa* L.) is a most important food grain because it is consumed by more than half of the world’s population for their sustainable livelihood (World Rice Production, 2019). It was domesticated in South East-South Asia and is now cultivated throughout the world. The japonica type was first originated and domesticated from a specific gene pool of *O. rufipogon* and indica type was subsequently developed from crosses between japonica rice and local wild rice (Huang et al., 2012). Two different models were proposed for the domestication of cultivated rice (*O. sativa*). One model proposed single domestication that *O. sativa* japonica type was initially originated from the progenitor *O. rufipogon* and acquired the domestication phenotypes then introgressed to other predomesticated populations giving indica and aus varieties (common origins). Other model proposed multiple domestication that japonica and indica type domesticated independently from the ancestral progenitor *O. rufipogon*. Some evidence support the models (independent origins) in which japonica, indica and aus were domesticated independently (Kovach et al., 2007; Civan and Brown 2018). Cultivated rice *O. sativa* can be divided into five groups based on genetics and culinary properties such as japonica (subdivided into tropical and temperate japonica), indica, aus and aromatic (Garris et al., 2005; Zhao et al., 2011; Civan et al., 2015). The indica genome appears to be least affected by gene-flow. Three independent origins for japonica, indica and aus were established (Civan and Brown 2018) based on genetic introgression analysis. Study on intron sequence and retrotransposon insertions suggest that at least two independent domestications of *O. sativa* from pre-differentiated gene pools of the *O. rufipogon* wild ancestor (Vaughan, 2003; Garris et al., 2005; Second, 1982; Londo et al., 2006) was happened and all of these calculations place the time to the recent common ancestor at more than 100,000 years ago (Vaughan, 2003). It is widely documented that domestication is not a single event, but rather a dynamic evolutionary process that occurs over time. Molecular phylogenetic analyses indicated that indica and japonica rice originated independently but other evidences support single domestication origin. Evolutionary history of rice domestication is still in debatable conditions. Some of the studies reported that progenitor wild rice (*O. rufipogon*) where from aus type rice was originated distributed from Brahmaputra valley through Bangladesh to the Odisha region in India. Japonica specific wild accession was most prevalent in Yangtze valley and Southern China. Although, indica specific wild rice variant not concentrated in any particular region but samples with highest proportions are identified in Indochina and the eastern part of the Indian subcontinent (Civan and Brown 2018). That means place of origin and domestication of rice is still in debatable conditions. Wild rice (*Oryza rufipogon* Griff.) is an immediate ancestral progenitor of cultivated rice (*O. sativa* L.) has red colour grain and most of the cultivated rice (*O. sativa* L.) has white grain. Grain colour varies from brown, red, black to white depending upon the accumulation of different flavonoid pigments. Wild rice (*O. rufipogon*) shows various pigmentation in the plant parts-black hull, red pericarp, purple awn, purple leaf margin. Most of the cultivated rice has lost colouration in these plant parts due to long term artificial selection and improvement (Sun et al., 2018).

Red pericarp is an inherent trait of wild rice *O. rufipogon*. White grain rice appears to be associated with domestication syndrome and is under strong selection pressure. The red pigment in rice pericarp is proanthocyanidin (condensed tannins), with important deterrent effects on pathogens and predators (Shirley, 1998). Rice domestication history reflects the trait selection for agronomically important characters such as seed shattering, plant architecture, yield and grain colour (Izawa et al 2009). Domesticated related genes in rice Sh4, qSH1, PLOG1, qSW5, GIF1 and Rc have been identified and cloned with FNP (Konishi et al., 2006; Sweeney et al., 2007; Wang et al., 2008). Domestication genes Bh4 (hull colour), PROG1 (tiller angle), sh4 (seed shattering), qSW5 (grain width), OsC1 (leaf sheath colour and apiculus colour), qSH1 (seed shattering), Waxy (grain quality), Rc (pericarp colour) have been characterized and identified (Huang et al., 2012). Other domestication genes GhD7, LABA1, Kala4, LG1 has also been analyzed through genetic analysis (Civan and Brown 2018). Most of the wild rice grain is red while cultivated rice grain is white (colourless) has been selected during domestication. Grain colour domestication genes Rc and Rd were responsible for rice pericarp pigmentation (Konishi et al. 2008; Sweeney et al. 2006; Sweeney et al. 2007). Red rice colour is attributed by two genes Rc and Rd. Loss-of-function mutation of Rc produces white grains in cultivated varieties. A 14-bp deletion was identified in the Rc gene of white pericarp cultivars. Maximum number of rice varieties (97%) with white pericarp carries a 14-bp deletion in the Rc gene which leads to a premature stop codon due to frameshift mutation and as a result nonfunctional *rc* allele is generated, which encodes truncated protein lacking bHLH domain and turns red pericarp into white pericarp. Remaining rice varieties (3%) with white pericarp, show a C-A point mutation in the exon 7 which is a null mutation and designated as Rc-s. It is a rare mutation in rice. It was demonstrated from the haplotype analysis of Rc gene, revealed that the 14-bp deletion arose from japonica and was introgressed into indica and aus, and the C-A mutation originated in the aus subpopulation and was not widely disseminated during domestication (Sweeney et al. 2007). Some of the findings suggested that the domestication process of white pericarp in aus is different from that in japonica and indica (Wang et al. 2016). Rc is expressed in both red and white pericarp varieties, but in white grained varieties, the Rc transcript is shortened due to the 14-bp deletion (Sweeney et al. 2006). White rice grain trait was spread into most rice varieties in the process of domestication (Sweeney et al., 2007). At present two genes are necessary to the red pigment synthesis in rice, Rc and Rd. Rc encodes bHLH transcription factor and Rd encodes dihydroflavonol-4-reductase (DFR) (Sweeney et al., 2006), Furukawa et al., 2007). Functional changes in Rc and Kala4 gene regulate the common biosynthesis pathway for the pigment formation which arose within a relatively evolutionary short period of time and produce distinct products (or trait), to produce diversity within plant species (Oikawa et al., 2015). Rc is closely associated with shattering and dormancy traits, thus selected against during domestication (Mbanjo et al., 2020). Rc-Rd genotypes produce red grain, while Rc-rd genotypes produce brown grain (Furukawa et al., 2006; Sweeney et al., 2006). Three alleles of Rc gene are Rc wild type, and mutant alleles rc and Rc-s. Mutant alleles rc, lacks 14 bp sequence of the wild type Rc allele, Rc-s is a point mutation C – A transversion leads to premature stop codon. Rice varieties carry rc allele produce white grain (colourless) while those of Rc-s produce a range of pericarp pigmentation. Accumulation of rice anthocyanins in the pericarp is a domestication related trait, which sheds light on the evolutionary history of rice (Duo et al., 2020). Most abundant anthocyanin is cyanidin 3-O-glucoside (Abdel-Aal et al. 2006) in the coloured rice. Red rice accumulates flavonoid proanthocyanins in the grain (Reddy et al. 1995). Three different types of regulatory genes such as transcription factor MYB, basic helix-loop-helix (bHLH) and WD40 regulates the anthocyanin biosynthesis in plants. Purple rice leaf colour is associated with OsPl gene MYB type factor and control the biosynthesis of anthocyanin and mutant version shows defect in stress response (Akhter et al., 2019). Natural selection favored red rice development not because of pericarp colour and nutritional value but probably because of its seed dormancy, which enhances survival of seeds and distributes germination over time (Gu et al., 2011). Red pericarp developed due to the accumulation of proanthocyanins and black rice contains anthocyanin, both are shared some identical biosynthetic genes (Reddy et al. 1995; Tomoyuki et al. 2002; Winkel-Shirley 2001). Rc and Rd have been projected to be the responsive genes of the red pericarp formation in rice grain. Functional alleles of Rc and Rd together develop red pericarp, functional allele of Rc alone can produce brown colour pericarp, and functional allele of Rd alone produces only white pericarp (Shao et al. 2011). It was demonstrated that two important domestication alleles which are unlinked rc (white pericarp) and the seed shattering allele, sh4 are located on Chromosome 4 and now estimated to have arisen only once in evolution and to have been introgressed across the indica–japonica subspecies, prevalent in the present day cultivated rice (Li et al., 2006).

Genetic loci such as Kala4/OsB2/Pb are reported to be indispensable for the synthesis of black (purple) pericarp in rice grain (Maeda et al. 2014; Rahman et al. 2013; Wang and Shu 2007). An insertional mutation was detected at the promoter region of Kala4 gene locus, which is a functional-gained mutation. As a result expression level of Kala4 gene significantly enhanced. Evidence suggests that this insertion was reported to be the original genetic change that gave birth to black rice (Oikawa et al. 2015). Several studies reported that the nucleotide diversity of Kala4 locus in cultivated rice varieties predicted that the insertion first took place in tropical japonica, and then spread to indica and subsequently temperate japonica through natural crossing and artificial selection of black rice trait (Oikawa et al. 2015). One QTL rg7.1 on chromosome 7 was characterized in red pericarp phenotype (Septiningsih et al., 2003), and analysis revealed that LOC_Os07g11020 was the causal gene for rg7.1, which encodes a bHLH transcription factor. Thus, LOC_Os07g11020 is the Rc gene. Red pericarp contains proanthocyanidin and is regulated by Rc (bHLH) with functional Rd allele, whereas brown pericarp is formed without Rd allele (Sweeney et al., 2006; Furukawa et al., 2007). White grain induced by mutant allele of Rc gene, a bHLH gene. QTL mapping approach has been utilized to identify gene associated to black pigmentation in rice grain (Tan et al., 2001; Dong et al., 2008; Matsuda et al., 2012; Korte and Farlow 2013; Xu et al., 2017; Yang et al., 2018).

There is no black grain colour in wild rice accessions (*O. rufipogon*) (www.gramene.org/). Therefore, it is predicted that black rice trait is recently acquired by the varieties (Oikawa et al., 2015), and incorporated during domestication or after domestication of the cultivated rice. Black was used by the ancient Chinese dynasty (Newman, 2004) and therefore designated as emperor’s rice or forbidden rice for its rarity and nutritional value. There is no clear genetic knowledge that how the black grain (or purple grain) trait arose and incorporated into the different rice types (indica, japonica, aus, aromatic) (Oikawa et al., 2015).

Origin of black rice and spread of the trait remains fully undiscovered. Black rice grain due to the accumulation of anthocyanin pigment under the control of Kala4 bHLH gene. Expression of Kala4 gene is possible due to rearrangement of promoter region. Two major genes Rc and Kala4 jointly upregulate the upstream gene of the flavonol biosynthesis gene-chalcone synthase, dihydroflavonon-4-reductase and downstream genes leucoanthocyanidin reductase and leucoanthocyanidin dioxygenase, to synthesize anthocyanin and proanthocyanidin pigment in the pericarp. Red grain colour is due to the deposition of proanthocyanidins in the pericarp and simultaneously oxidative polymerization. Black rice accumulates anthocyanin pigment in the pericarp (Rahman et al., 2013, Goufo and Trindade 2014). Anthocyanin biosynthesis and transcriptional regulation has been studied in many plant species and maize (Shih et al., 2008). Regulatory genes R/B and C1/P1 gene families in maize and orthologs in other plants demonstrated these encode bHLH (basic helix-loop-helix) type and R2R3-MYB type transcription factors respectively (Chandler et al., 1989; Cone et al., 1993). All three factors act as ternary complex as MBW (MYB-bHLH-WD40) (Hichri et al., 2011; Petroni and Tonelli 2011). Purple leaf colour regulated by (Pl) gene with three allelic variation Pl^i^, Pl^j^, Pl^w^. Same locus revealed two candidate R genes, OSB1 and OSB2 (Sakamoto et al., 2001), activate the biosynthesis of anthocyanin but not yet confirmed experimentally. There are few reports about the breeding and inheritance of black grain trait in the progeny populations. Black rice introgression lines have been developed by crossing and backcrossing ‘Hong Xie Nuo’ (black rice cultivar) with ‘Koshihikari’(white grain cultivar) (Maeda et al 2014). Breeding study of a cross between white cultivar ‘Koshihikari’ and black rice ‘Hong-Xie-Neo’ reported that black pigmentation controlled by mainly three genes such as Kala1, Kala3 and Kala4, during the black trait inheritance in the NIL lines (Maeda et al., 2014). Speculated from the crossing, that white grain was converted to black grain in the NIL lines (Near isogenic line) and demonstrated that the allele Kala1 and Kala3, those encode DFR and R2R3-Myb transcription factor respectively are important for the synthesis of anthocyanin in the pericarp (Maeda et al., 2014). In this NIL line, one major allele Kala4 was indentified and characterized. The Kala4 was considered as a responsible allele for origin of black rice. It was located on chromosome 4 at sequence position Os04g0557500, encodes a bHLH transcription factor, homolog of maize R/B gene. It was previously reported to be OSB2 gene (Sakamoto et al., 2001). Structural change and rearrangement of the promoter region of Kala4 allele induced ectopic expression of bHLH transcription protein factor which subsequently activate the synthesis of anthocyanin in rice pericarp that initiate the birth of black rice. Small genomic segment containing Kala4 locus, originated from tropical japonica was introgressed into the indica subspecies through repeated natural out crossing and selection (Oikawa et al., 2015). It was reported that neo-functionalization of Kala4 allele, ultimately give birth of black rice in the genetic background of tropical japonica and spread to other subspecies rice. Black pigmentations in rice pericarp are associated with the three gene loci, such as Kala1, Kala3 and Kala4 (Maeda et al 2014). Furthermore, Kala1 has been found to encode the DFR enzyme and mapped to chromosome 1 while Kala3 and Kala4 correspond to bHLH domain-containing and MYB domaincontaining transcription factors respectively. Three loci reportedly acted as a major determinant of pigment formation in black rice pericarp (Maeda et al 2014).

Previous studies showed that the 11.0 kb insertion is necessary to make positive changes in the Kala4 promoter region to express the black trait. Structural change in the promoter of Kala4 confers the black rice pericarp trait in rice varieties. Kala4 gene both with and without the LINE1 insertion were observed in rice landraces. It is not the reason for the ectopic expression of Kala4 gene to produce black rice pericarp. But LINE1 insertion makes the genomic region instable near the Kala4 region. That helps to generate duplication of Kala4 and subsequently promote ectopic expression. A part of the LINE1 retrotransposon element is duplicated during the rearrangement of promoter region of Kala4 allele through recombination events. Overall, 11.0 kb insertion following rearrangement *via* recombination was the key change in the promoter region which makes the Kala4 promoter functional and expressed for black colour pigment formation (Oikawa et al., 2015). Transposon is acted as modulators of gene expression during domestication of crop species (Studer et al., 2011). In case of black rice variety ‘Hong Xie Nuo’ Kala4 promoter possesses a duplication region of size 4.679 kb incorporating exon1, intron1, exon2 and also part of intron2. Besides the partial duplication,, the Kala4 promoter region in black rice variety (‘Hong Xie Nuo’) has an extra insertion of 11.02 kb genome segment taking from −83 upstream of the Kala4 gene (Oikawa et al., 2015). In the course of its evolution black rice possessed red pericarp.

Genetic analysis demonstrated that two genes PURPLE PERICARP B (Pb) located on chromosome 4 and PURPLE PERICARP A (Pp) located on chromosome 1, were required for black pigmentation. In addition to that a region (P) on chromosome 3, is also necessary for black grain formation. Three loci are needed for the synthesis of black pericarp, namely *Kala*1, *Kala*3 and *Kala*4. Grains are black if both the genes *Pb* and *Pp* are present, brown colour if *Pb* present while white grain if only *Pp* present. Three chromosomal loci *Kala*1, *Kala*3 and *Kala*4 are associated with the allelic loci (Pp) Rd; P, and Pb respectively (Maeda et al., 2014). Three allelic loci are linked to the possible genes, Rd/DFR, P/MYB and Pb/bHLH, which encodes dihydrofolate reductase (Furukawa et al., 2006), MYB transcription factor (Saito et al., 2004; Gao et al., 2011), and bHLH (basic Helix-Loop-Helix) transcription regulator respectively (Sakamoto et al., 2001). In red rice, Rd interacts with Rc in chromosome 7 to produce red grain. The chromosomal region of Kala 1 contains the Rd locus, which has been characterized as DFR gene (Furukawa et al., 2006). In red rice the Rd gene interacts with the Rc gene on chromosome 7, to produce red grains (Furukawa et al., 2006). Thus it appears likely that Kala 1 is Rd/DFR and Kala1 interacts with Kala4 in the production of coloured grain. Purple leaf (Pl) colour is controlled by three alleles these are *Pl^i^, Pl^w^*, and *Pl^j^*. Among these three alleles, *Pl^w^* confers pigmented pericarp (Sakamoto et al., 2001). The *Pl* gene encodes bHLH transcription factor. It is suggested that Kala1 and Kala4 are necessary for anthocyanin synthesis. The MYB gene functions as a tissue-specific gene regulator. As for example, the colourless like regulatory gene 1 (OsC1) on chromosome 6, encodes a MYB family transcription factor, responsible for the synthesis of anthocyanin in the apiculus and sheath (Saito et al., 2004; Gao et al., 2011). There are several report regarding the biosynthesis of anthocyanin and other pigments in rice grain (Gu et al., 2005; Deng et al., 2013; Pintha et al., 2015; Chen et al.,2016; Oikawa et al. 2015; Gu et al., 2011; Visioli, 2016; Zheng et al. 2019).

Colour intensity variation was observed in the black rice grains, which imply that the pigmentation trait is under polygenic control (many genes) and many genes are still unidentified (Ham et al., 2015). Several genes are identified and characterized for the trait of black pericarp in rice (Sakamoto et al., 2001, Rahman et al., 2013; Ham et al., 2015, Sakulsingharoj et al., 2014, 2016). The Pl locus carries two genes OSB1 and OSB2, each of which encodes a bHLH transcription factors (Sakamoto et al., 2001). Dominant complementary genes Pb (synonym Prp-b) and Pp (synonym Prp-a), located on chromosome 4 and 1 respectively determines the black pericarp colour in rice (Rahman et al., 2013). Pb for brown pericarp and Pp for purple pericarp, depends on the copy number of the gene (Rahman et al., 2013). Pb alone produces brown grain in absence of Pp allele. Lacking Pb, Pp forms white colour. Recently three genetic loci have been identified namely Kala1 (chromosome 1), Kala3 (chromosome 3), and Kala4 (chromosome 4) to produce black pigment in the rice pericarp (Maeda et al., 2014). Some study suggested that Kala4 is synonymous with Pb and Kala1 with Pp. Kala4 encodes a bHLH transcription factor protein and corresponds to OSB2 gene. The OSB2 regulates several genes which are encoding enzymes involved in anthocyanin biosynthesis, including F3H, DFR and ANS (Sakulsingharoj et al., 2014). Some study provides information that Kala1 locus include Rd region, located on chromosome 1, Kala3 is synonymous to MYB3, located on chromosome 3 (Maeda et al., 2014) and Kala4 is synonymous of bHLH transcription factor present on chromosome 4. Results of whole genome sequencing along with transcriptomic analysis in black rice varieties identified several genes involving in anthocyanin biosynthesis these are UGT, ANS1, F3H, and DFR (Oh et al. 2018). Anthocyanin biosynthesis in rice plant is responsive to abiotic stresses, such as drought, high salt and ABA, and also involved in biotic stress such as disease resistance (Gandikota et al. 2001; Ithal and Reddy 2004).

Black rice is considered as a panacea or super food because of its high nutritive value, curative effect coupled with beneficial properties of flavonoids that not only act as an antioxidant and anti-inflammatory but has also been shown to anti-carcinogenic properties. There are more than 200 types of black rice varieties in the world. Black rice grain has higher commercial value due to the presence of flavonoid compound anthocyanins that has antioxidant activity and is beneficial for human health. Black rice traits have been preferred by breeders and consumers for quite a long time and the cultivation process have been improved (Ito and Lacerda 2019). Market value of pigmented rice is potentially quite high due to growing consumer preference for nutritious foods (Islam et al., 2018). Looking forward, there is a tremendous opportunity for breeding programs to develop productive pigmented rice varieties (Voss-Fels et al., 2019). To meet up the market demands, several breeding approaches were initiated to develop new superior black rice variety taking local black rice as donor parent. A number of black rice lines developed through crossing between black rice ‘Okunomurasaki’ and white rice Koshihikari (Maeda et al., 2014). Riceberry, a newly developed black rice by crossing between aromatic Jasmine rice and local non-glutinous purple variety (Waiyawuththanapoom et al., 2015). The red rice ‘Rubi’ and black rice ‘Onix’ have been released in Brazil (Wickert et al., 2014), developed through breeding hybridization. Pigmented rice varieties are gaining popularity among consumers due to nutritional health benefits and market demands is expected to rise.

It was reported that Kala4 introgressed multiple times from japonica to indica rice types during domestication (Oikawa et al., 2015). The R2R3-MYB (Os06g0205100) functioning as a activator of DFR and ANS gene (Saitoh et al., 2004); Rachasima et al., 2017; Sun et al., 2018). Dihydroflavonol-4-reductase (Os01g0633500) involved in anthocyanin biosynthesis. Several authors have tried to make interrelationship between the genes and traits production leading to black pigmentation in rice (Sakulsingharoj et al., 2014; Rachasima et al., 2017; Sun et al., 2018; Lachagari et al., 2019; Mbanjo et al., 2020). There are a number of genes related to the black rice pigmentation in rice have already been detected and characterized. But there is still a possibility that some additional genes and allelic variants thereof remain to be exposed (Mbanjo et al., 2020). QTL mapping approach has been utilized to identify gene associated to black pigmentation in rice grain (Tan et al., 2001; Dong et al., 2008; Matsuda et al., 2012; Korte and Farlow 2013; Xu et al., 2017; Yang et al., 2018). Transcriptomics approaches have also been studied to elucidate the differential expression of the regulators and related genes to flavonoid biosynthesis in pigmented and non-pigmented rice grain (Lim and Ha, 2013; Oh et al., 2018). Sun et al (2018) studied the flavonoid synthesis pathway related genes and observed that these are under the control of ternary MYB-bHLH-WD40 transcriptional complex (Xu et al., 2013; Zhang et al., 2018). RNA-seq technique applied to identify the pigment biosynthesis pathway associated genes in rice (Kim et al., 2018). The bHLH transcription factor encoded by Rc allele activates the gene to synthesize dihydroflavonol-4-reductase, which influence the accumulation of red pigment in rice pericarp.

There is no report about the development of black rice through conventional breeding methods in the history of rice research by combining non-black rice varieties. Therefore, present study report in the first time to the rice genetics and breeding that black rice was developed through interspecific hybridization between rice cultivars (Badshabhog x *O. rufipogon*), (Ranjit x *O. rufipogon*) and (Chenga x *O. rufipogon*). Black rice was created by wide crossing in the genetic background of cultivated rice *O. sativa* through introgression of the black rice gene associated genomic segment into the pre-breeding lines which was inherited in the successive generations. It explores the genetic basis of origin of black rice and suggests a new model about the independent origin of black rice in the genetic background of indica rice variety and also provides information about the genetic rearrangement event during black domestication history.

## Materials and methods

### Plant material

Three rice [*Oryza sativa* L.) cultivars namely Badshabhog, Chenga, Ranjit and one wild rice genotype [*Oryza rufipogon* Griff.] were used for the present interspcific hybridization. Cultivar Badshabhog is aromatic with white grain, Chenga is non-aromatic with red grain, and Ranjit is a HYV non-aromatic with white grain. Wild rice genotype (*O. rufipogon*) is collected from local area Raiganj, West Bengal, India (Fig. 1); conserved and maintained in the Plant Genetics & Molecular Breeding Laboratory, Department of Botany; University of North Bengal, India.

**Figure 1.**
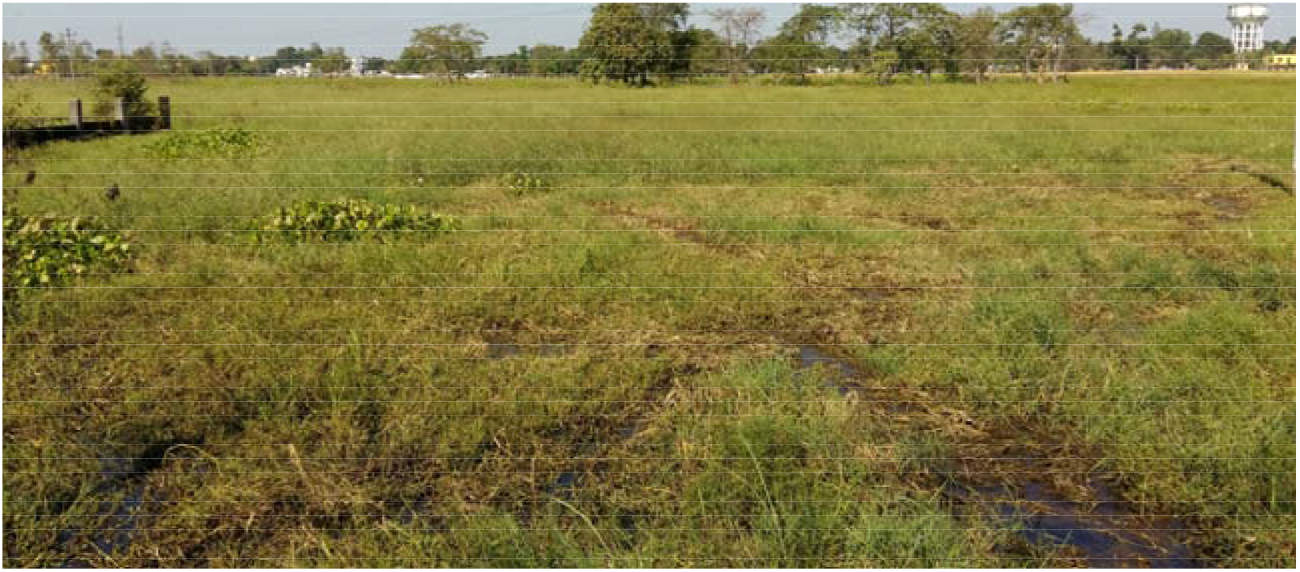
Natural growing habitat of wild rice (*Oryza rufipogon* Griff.) at Raiganj, West Bengal, India, used in this interspecific hybridization for the development of black rice.

### Hybridization and artificial pollination

Pre-breeding lines were developed by artificial pollination and maintained for the present work. Hybridization was done in the year 2016 based on standard protocol of artificial pollination (Sleper and Poehlman 2007; Roy, 2017; Roy and Reddy 2017; Roy and Shil 2020) for the development of breeding lines (Badshabhog x *O. rufipogon*; Ranjit x *O. rufipogon*) and Chenga x *O. rufipogon* in the year 2017. After 25 days of pollination F1 seeds were collected from the parent plant (November 2016) and then subsequently maintained the lines for the development of F5 populations. Agro-morphological traits were recorded according to SES Bioversity-IRRI DUS test protocol (2014) for characteristics evaluation of the breeding lines. Observations like plant height, panicle length, grain per panicle, grain length, grain breadth, and 1000 grain weight were recorded.

### Physicochemical properties and sensory based aroma test

Alkali spreading value (ASV) (in a scale of 1 to 7) was measured based on standard protocol (Little et al., 1958). A low ASV corresponds to a high gelatinization temperature (GT), conversely, a high ASV indicates a low GT. Sensory based aroma (in a scale of 0 to 3) was assessed using standard method (Sood and Siddiq, 1978).

## Results

### Breeding lines developed

Pre-breeding lines were developed and maintained for genetic improvement of the cultivated varieties before release to the farmers’ field. Progeny F5 lines (Badshabhog x *O. rufipogon*; Ranjit x *O. rufipogon*)), and progeny F4 lines (Chenga x *O. rufipogon*) were maintained and evaluated at the field conditions. Pre-breeding lines were evaluated for the introgression of agronomically important traits (gene/QTLs) into the cultivars at F_2_ segregation populations (Ranjit x *O. rufipogon*; Badshabhog x *O. rufipogon*; Chenga x *O. rufipogon*). Agro-morphological characteristics of each of the pre-breeding lines were evaluated for any unique traits development starting from F_2_ segregating populations and onwards F5 for (Ranjit x *O. rufipogon*; Badshabhog x *O. rufipogon*) and F4 for (Chenga x *O. rufipogon*) at present Kharif 2020. All the breeding lines were maintained for genetic evaluation purpose at the field level based on SES IRRI test protocol (Fig. 2).

**Figure 2.**
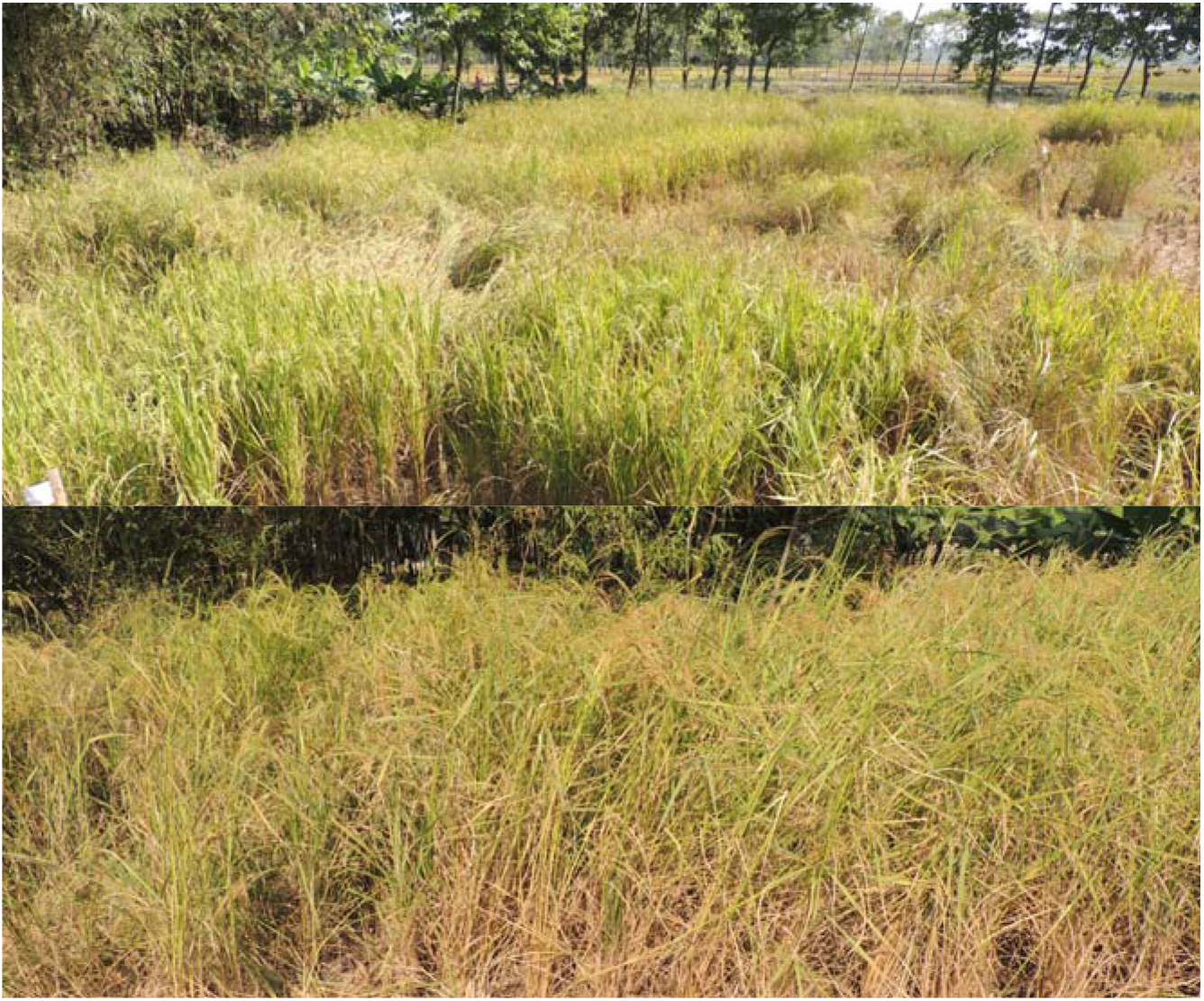
Pre-breeding lines were evaluated at field conditions at F5 (Badshabhog x *O. rufipogon*); (Ranjit x *O. rufipogon*) and at F4 populations level (Chenga x *O. rufipogon*) for trait expression and plant architecture observations (Kharif crop 2020).

### Agro-morphological variability parameters

Statistical analysis Anova was performed and analysis revealed that there is high significant difference between the genotypes in respect to fifteen (15) traits studied (Table 1–3) comprising plant height (PH), flag leaf length (FLL), flag leaf width (FLB), panicle length (PnL), grain per panicle (Gr/Pn), grain length (GL), grain breadth (GB), 1000 grain weight (GrWt), heading date (HD), maturity time in days (MT), active tillering number (Till), awn length (Awn), shattering trait, pericarp colour and aroma. It suggested that there were inherent genetic differences among the breeding lines (genotypes) in respect to the morphological traits considered during the analysis. High variability of breeding lines will increase the probability of producing desirable introgression and recombinants in successive generations.

**Table 1.** Agro-morphological traits of pre-breeding (Badshabhog x *O. rufipogon*) lines at F5 generation were evaluated based on DUS Test protocol.

| PLANT | PH(cm) | PnL(cm) | FLL(cm) | FLB(mm) | GL(mm) | GB(mm) | Gr/Pn | GrWt | Till | HD | MT | Aroma | AWN (mm) | Pericarp colour | Shattering traits |
| --- | --- | --- | --- | --- | --- | --- | --- | --- | --- | --- | --- | --- | --- | --- | --- |
| Local Black Rice (control) | 155.50±0.55 | 26.50±0.35 | 47.00±0.35 | 15.00±0.14 | 8.75±0.02 | 3.12±0.01 | 70.00±4.85 | 24.00±0.08 | 09.00±0.65 | 110.00±0.00 | 150.00±0.00 | 3 | Nil | Black | Nsh |
| Badshabhog Parent 1 | 145.00±0.51 | 28.00±0.26 | 35.00±0.38 | 11.19±0.12 | 6.13±0.02 | 1.98±0.02 | 362.00±9.58 | 10.60±0.05 | 12.00±0.59 | 110.00±0.00 | 145.00±0.00 | 3 | Nil | White | Nsh |
| BWL1 | 155.45±0.55 | 27.50±0.31 | 37.50±0.30 | 11.50±0.09 | 9.51±0.03 | 2.68±0.01 | 48.21±2.05 | 18.00±0.09 | 22.25±0.73 | 110.00±0.00 | 145.00±0.00 | 3 | 20.00±2.18 | Red | SmSh |
| *O.rufipogon* Parent 2 | 160.00±1.58 | 18.00±0.35 | 17.50±0.59 | 7.24±0.19 | 8.52±0.02 | 2.11±0.016 | 38.00±1.93 | 17.20±0.06 | 95.00±3.63 | 125.00±0.00 | 150.00±0.00 | 0 | 80.00±3.15 | Red | Sh |
| BWL2 | 158.02±0.65 | 24.50±0.39 | 29.61±0.39 | 12.21±0.08 | 5.60±0.02 | 2.35±0.009 | 105.30±6.95 | 11.00±0.05 | 19.10±0.84 | 110.00±0.00 | 145.00±0.00 | 3 | Nil | Red | Nsh |
| BWL3 | 165.31±0.66 | 21.70±0.42 | 26.30±0.48 | 10.14±0.07 | 6.97±0.03 | 2.68±0.02 | 107.33±7.10 | 10.00±0.06 | 29.09±0.95 | 110.00±0.00 | 145.00±0.00 | 0 | 31.00±1.89 | Red | SmSh |
| BWL4 | 152.22±0.68 | 29.50±0.31 | 39.30±0.53 | 12.25±0.10 | 8.06±0.02 | 2.95±0.02 | 185.45±9.15 | 27.00±0.07 | 28.07±1.06 | 110.00±0.00 | 145.00±0.00 | 0 | Nil | White | Nsh |
| BWL5 | 161.43±0.55 | 30.60±0.42 | 41.20±0.31 | 11.30±0.09 | 7.45±0.02 | 2.43±0.01 | 178.10±8.84 | 10.00±0.06 | 32.20±0.89 | 110.00±0.00 | 145.00±0.00 | 0 | Nil | White | Nsh |
| BWL6 | 148.10±0.61 | 19.20±0.30 | 25.21±0.40 | 10.20±0.08 | 7.40±0.04 | 2.56±0.02 | 75.45±4.10 | 19.00±0.08 | 22.70±1.10 | 110.00±0.00 | 145.00±0.00 | 0 | Nil | Red | Nsh |
| BWL7 | 160.08±0.75 | 18.00±0.43 | 39.10±0.40 | 10.30±0.07 | 9.02±0.04 | 2.88±0.02 | 57.74±2.88 | 20.00±0.09 | 12.96±0.89 | 110.00±0.00 | 145.00±0.00 | 3 | Nil | Black | Nsh |
| Local Black Rice (Control) | 155.50±0.55 | 26.50±0.35 | 47.00±0.35 | 15.00±0.14 | 8.75±0.02 | 3.12±0.01 | 70.00±4.85 | 24.00±0.08 | 9.00±0.65 | 110.00±0.00 | 150.00±0.00 | 3 | Nil | Black | Nsh |
| Badshabhog | 145.00±0.51 | 28.00±0.26 | 35.00±0.38 | 11.19±0.12 | 6.13±0.02 | 1.98±0.02 | 362.00±9.58 | 10.60±0.05 | 12.00±0.59 | 110.00±0.00 | 145.00±0.00 | 3 | Nil | White | Nsh |
| BWL8 | 155.45±0.55 | 27.50±0.31 | 37.50±0.30 | 11.50±0.09 | 9.51±0.03 | 2.68±0.01 | 48.21±2.05 | 18.00±0.09 | 22.25±0.73 | 110.00±0.00 | 145.00±0.00 | 3 | 20.00±2.18 | Red | SmSh |
| *O.rufipogon* | 160.00±1.58 | 18.00±0.35 | 17.50±0.59 | 7.24±0.19 | 8.52±0.02 | 2.11±0.016 | 38.00±1.93 | 17.20±0.06 | 95.00±3.63 | 125.00±0.00 | 150.00±0.00 | 0 | 80.00±3.15 | Red | Sh |
| BWL9 | 145.17±0.61 | 26.40±0.53 | 36.20±0.31 | 12.10±0.11 | 8.65±0.03 | 2.65±0.02 | 85.12±3.56 | 18.00±0.07 | 32.04±0.79 | 110.00±0.00 | 150.00±0.00 | 3 | Nil | Black | Nsh |
| BWL10 | 148.23±0.60 | 26.50±0.62 | 35.50±0.34 | 11.50±0.09 | 8.79±0.05 | 2.78±0.03 | 62.24±2.86 | 20.00±0.08 | 28.10±1.12 | 110.00±0.00 | 150.00±0.00 | 3 | Nil | Black | Nsh |
| BWL11 | 150.20±0.58 | 25.60±0.58 | 34.20±0.48 | 10.50±0.08 | 7.65±0.04 | 2.61±0.02 | 49.80±2.10 | 22.00±0.09 | 29.09±1.10 | 110.00±0.00 | 150.00±0.00 | 3 | Nil | Black | Nsh |
| BWL12 | 152.18±0.65 | 25.80±0.55 | 35.30±0.39 | 11.20±0.08 | 9.05±0.04 | 2.51±0.02 | 61.16±5.12 | 20.00±0.09 | 26.07±0.98 | 110.00±0.00 | 150.00±0.00 | 3 | 25.50±0.95 | Black | Nsh |
| BWL13 | 145.20±0.60 | 19.22±0.43 | 38.50±0.47 | 10.60±0.07 | 8.10±0.04 | 2.20±0.03 | 71.21±4.48 | 20.00±0.07 | 24.41±1.18 | 110.00±0.00 | 150.00±0.00 | 2 | Nil | White | Nsh |
| BWL14 | 147.38±0.68 | 26.30±0.41 | 39.50±0.45 | 11.20±0.07 | 7.10±0.03 | 2.30±0.02 | 79.32±5.12 | 12.00±0.06 | 28.14±1.20 | 110.00±0.00 | 150.00±0.00 | 3 | 28.90±0.86 | White | Nsh |
| BWL15 | 148.46±0.65 | 25.52±0.42 | 37.20±0.34 | 12.10±0.08 | 8.43±0.03 | 2.88±0.02 | 73.13±4.23 | 20.00±0.08 | 29.22±1.18 | 110.00±0.00 | 150.00±0.00 | 3 | Nil | Black | Nsh |
| BWL16 | 150.11±0.61 | 24.00±0.36 | 36.50±0.23 | 13.10±0.08 | 6.02±0.04 | 2.20±0.02 | 89.00±4.56 | 12.00±0.09 | 27.21±0.99 | 110.00±0.00 | 150.00±0.00 | 2 | Nil | White | Nsh |
| BWL17 | 139.32±0.56 | 23.51±0.38 | 34.50±0.25 | 11.00±0.07 | 6.58±0.04 | 2.59±0.02 | 165.31±6.98 | 9.00±0.09 | 28.23±1.49 | 110.00±0.00 | 150.00±0.00 | 3 | Nil | White | Nsh |
| BWL18 Short PH | 70.00.10±0.34 | 8.51±0.22 | 12.30±0.10 | 6.10±0.03 | 8.45±0.02 | 2.52±0.009 | 7.08±0.45 | 18.00±0.05 | 29.27±0.55 | 110.00±0.00 | 150.00±0.00 | 3 | 22.50±0.98 | White | SmSh |
| BWL19 (Bi colour seed) | 140.33±0.51 | 27.51±0.18 | 33.50±0.21 | 11.50±0.09 | 8.02±0.03 | 2.56±0.02 | 112.12±7.88 | 18.00±0.07 | 24.29±1.56 | 110.00±0.00 | 150.00±0.00 | 3 | Nil | White | Nsh |
| Badshabhog | 145.00±0.51 | 28.00±0.26 | 35.00±0.38 | 11.19±0.12 | 6.13±0.02 | 1.98±0.02 | 362.00±9.58 | 10.60±0.05 | 12.00±0.59 | 110.00±0.00 | 145.00±0.00 | 3 | Nil | White | Nsh |
| BWL20 | 204.51±0.71 | 23.50±0.39 | 32.85±0.24 | 15.10±0.08 | 8.12±0.03 | 2.23±0.02 | 125.45±8.84 | 22.15±0.07 | 27.25±1.83 | 110.00±0.00 | 150.00±0.00 | 0 | 72.59±0.95 | Red | Sh |
| BWL21 Shortest PH | 60.12±0.28 | 10.83±0.25 | 12.23±0.11 | 5.11±0.02 | 6.05±0.02 | 2.12±0.03 | 11.79±1.02 | 6.29±0.03 | 17.33±0.95 | 110.00±0.00 | 150.00±0.00 | 0 | 24.85±0.86 | White | Sh |
| BWL22 | 152.44±0.63 | 28.13±0.32 | 43.21±0.48 | 12.20±0.09 | 7.12±0.03 | 1.76±0.02 | 92.55±7.56 | 15.04±0.09 | 29.21±1.28 | 110.00±0.00 | 150.00±0.00 | 3 | 3.26±0.12 | White | SmSh |
| BWL23 | 170.21±0.69 | 31.50±0.39 | 46.87±0.31 | 21.52±0.07 | 7.75±0.02 | 2.57±0.02 | 278.54±12.34 | 17.51±0.06 | 26.85±1.98 | 110.00±0.00 | 150.00±0.00 | 0 | Nil | White | Nsh |
| *O.rufipogon* | 160.00±1.58 | 18.00±0.35 | 17.50±0.59 | 7.24±0.19 | 8.52±0.02 | 2.11±0.016 | 38.00±1.93 | 17.20±0.06 | 95.00±3.63 | 125.00±0.00 | 150.00±0.00 | 0 | 80.00±3.15 | Red | Sh |

**Table 2.**
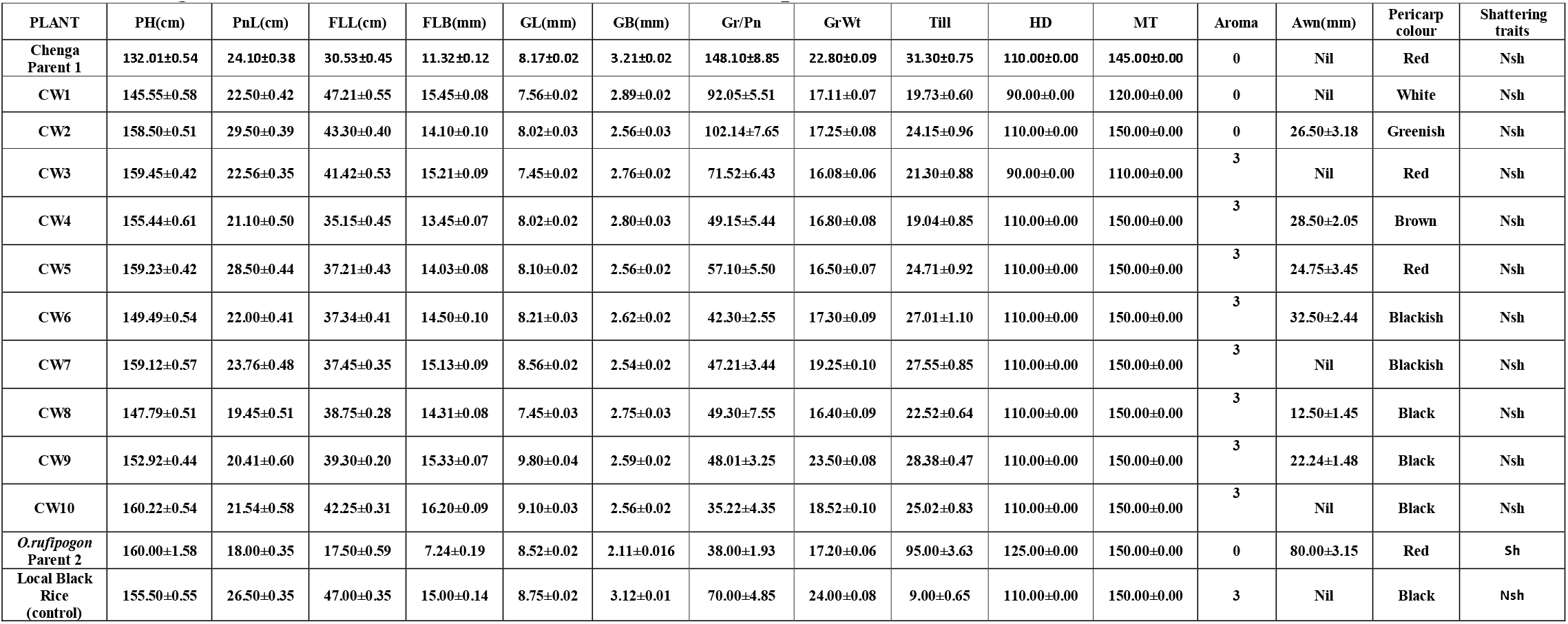
Agro-morphological traits of pre-breeding (Chenga x *O. rufipogon*) lines at F4 generation were evaluated based on DUS Test protocol.

**Table 3.** Agro-morphological traits of pre-breeding (Ranjit x *O. rufipogon*) lines at F5 generation were evaluated based on DUS Test protocol.

| PLANT | PH(cm) | PnL(cm) | FLL(cm) | FLB(mm) | GL(mm) | GB(mm) | Gr/Pn | GrWt (gm) | Till | HD | MT | Aroma | Awn(mm) | Pericarp colour | Shattering traits |
| --- | --- | --- | --- | --- | --- | --- | --- | --- | --- | --- | --- | --- | --- | --- | --- |
| Ranjit Parent 1 | 111.50±0.40 | 21.75±0.22 | 23.14±0.43 | 16.00±0.13 | 7.50±0.02 | 2.97±0.01 | 213.57±2.37 | 20.80±0.09 | 16.20±0.68 | 85.00±0.00 | 125.00±0.00 | 0 | Nil | White | Nsh |
| RRL1 | 145.52±0.55 | 26.40±0.24 | 32.50±0.51 | 15.70±0.07 | 7.25±0.02 | 2.54±0.01 | 95.60±6.11 | 17.20±0.07 | 21.12±0.98 | 90.00±0.00 | 130.00±0.00 | 0 | Nil | Red | Nsh |
| RRL2 | 165.00±0.61 | 31.50±0.36 | 38.74±0.56 | 16.12±0.09 | 5.21±0.02 | 2.56±0.02 | 186.10±10.10 | 12.19±0.08 | 27.10±0.77 | 110.00±0.00 | 150.00±0.00 | 0 | 3.10±0.50 | White | Nsh |
| RRL3 | 155.23±0.48 | 30.25±0.59 | 37.34±0.45 | 15.50±0.06 | 7.95±0.02 | 2.56±0.02 | 95.25±5.88 | 18.22±0.08 | 28.09±1.38 | 110.00±0.00 | 150.00±0.00 | 0 | Nil | White | Nsh |
| RRL4 | 154.25±0.53 | 30.52±0.22 | 39.45±0.61 | 14.58±0.08 | 8.20±0.03 | 2.48±0.01 | 194.30±9.88 | 19.40±0.06 | 18.30±1.65 | 90.00±0.00 | 130.00±0.00 | 0 | Nil | Red | Nsh |
| RRL5 | 169.23±0.55 | 37.50±0.58 | 46.50±0.73 | 15.50±0.10 | 7.05±0.02 | 2.21±0.01 | 94.60±5.10 | 14.21±0.06 | 26.12±1.38 | 110.00±0.00 | 150.00±0.00 | 0 | Nil | Red | Sh |
| RRL6 | 165.24±0.59 | 21.30±0.24 | 33.25±0.61 | 14.25±0.12 | 7.89±0.02 | 2.56±0.02 | 96.70±7.88 | 18.14±0.08 | 21.07±0.99 | 110.00±0.00 | 150.00±0.00 | 0 | 10.20±2.59 | Red | Nsh |
| RRL7 | 142.50±0.41 | 22.45±0.41 | 32.50±0.42 | 14.00±0.11 | 8.10±0.02 | 2.20±0.01 | 75.30±6.85 | 17.11±0.07 | 13.30±1.23 | 90.00±0.00 | 130.00±0.00 | 0 | Nil | White | Nsh |
| RRL8 | 155.50±0.58 | 26.80±0.34 | 35.60±0.36 | 15.40±0.09 | 8.27±0.03 | 2.56±0.02 | 96.50±6.51 | 18.08±0.08 | 14.50±1.45 | 110.00±0.00 | 150.00±0.00 | 0 | 25.55±8.88 | Red | Nsh |
| RRL9 | 145.23±0.59 | 28.45±0.56 | 38.54±0.57 | 15.45±0.12 | 8.10±0.02 | 2.50±0.020 | 203.25±10.21 | 14.30±0.07 | 16.08±1.65 | 110.00±0.00 | 150.00±0.00 | 0 | Nil | White | Nsh |
| RRL10 | 128.00±0.60 | 24.25±0.48 | 25.60±0.49 | 15.24±0.13 | 7.08±0.02 | 2.53±0.01 | 165.55±6.05 | 17.10±0.08 | 12.10±1.69 | 90.00±0.00 | 130.00±0.00 | 0 | Nil | White | Nsh |
| *O. rufipogon* Parent 2 | 160.00±1.17 | 18.00±0.31 | 17.50±0.59 | 7.24±0.19 | 8.52±0.02 | 2.11±0.01 | 38.00±1.93 | 17.00±0.06 | 95.00±3.63 | 125.00±0.00 | 150.00±0.00 | 0 | 80.20±15.55 | Red | Sh |

### Black rice selection at F2 segregation population and inheritance of the black rice gene

It was unexpected to observe the black rice grain in the F2 segregating populations of pre-breeding lines (Badshabhog x *O. rufipogon*) and (Chenga x *O. rufipogon*). We detected and identified some of the individual plant with black grain, while we are taking the agro-morphological measurement at the harvesting time. Frequency was not so high; we found few plants with black grains from large populations (more than 8000 plants of each population) of wide crosses (Fig. 2). Some black rice grain is with aroma and according to the standard sensory based aroma test, it was at index level 3 (Table 1–3). Agro-morphological diversity was observed and analyzed about fifteen traits (Table 1–3). In the breeding lines (Badshabhog x *O. rufipogon*) variability was high in plant height trait (PH) ranges from 60 cm to 204 cm; shortest panicle length was recorded in the breeding line BWL18 only 8.51 cm and longest in breeding line BWL23 was 31.50 cm, grain length varies 5.60 mm to 9.51 mm in the lines BWL2 and BWL8 respectively, 1000 grain weight (GrWt) varies from 6.29g to 27.00g (Table 1 and Fig. 3–5). Some of the breeding lines show transgressive segregation in respect to 1000 GrWt, grain length, and panicle length (Table 1). Most interesting trait in this breeding line was development of black rice (black pericarp). Out of 23 BWL lines (Badshabhog x *O. rufipogon*) we selected for the present investigation, six breeding lines (BWL7, BWL9 - BWL12, and BWL15) were with new trait, black rice grain (Table 1 and Fig. 3–5). New trait, black rice grain formation is the unique invention in this prebreeding experiment. Not only black grain rice we detected from the breeding populations (Table 1, Fig. 3–5), but also detected grains with colour variation like-brown, red and white. It is supporting the view that grain colour is polygenetic inheritance controlled by many genes, or quantitative trait loci (QTL). Similar results were also observed in the breeding lines of the cross (Chenga x *O. rufipogon*) (CW). Grain colour variation also observed ranging from red, white, black, and greenish in this CW lines (Table 2, Fig. 6). We selected best performing three aromatic black rice lines from this cross (Table 2 and Fig. 6). We identified black rice from this second cross (Chenga x *O. rufipogon*), support the findings of first cross (Badshabhog x *O. rufipogon*) that black rice grain colour really has been developed through the interspecific crossing between *O. sativa* x *O. rufipogon*. Origin of black rice has been happened through the out crossing and introgression. Genetically it is quite viable and possible to create rearrangement in the breeding lines to induce the biosynthetic pathway associated with the anthocyanin pigment accumulation in the rice pericarp leading to black rice development. In the third cross between (Ranjit x *O. rufipogon*), black rice variant was not observed (Table 3 and Fig. 7). Ranjit is white grain rice HYV and wild rice (*O. rufipogon*) with red grain. In this third cross, we have detected only red and white grain, without any black rice generation. Supporting the proposition that black rice development needs some specific type of genotype. So that, it can provide cellular environment to reconstruct the genetic component in such a way, that it will induce genes and transcription modulator to start pigment biosynthesis mainly anthocyanin to give birth of black rice in the genetic background of indica rice varieties. Anthocyanin biosynthesis pathway associated genes and genetic components may be interact and complement each other to create functional gene cassette for the production of black grain pericarp. Brown pericarp form when only Rc is functional without Rd allele. All white grain is due to mutant *rc* allele. The Kala4 was considered as a responsible allele for origin of black rice.

**Figure 3.**
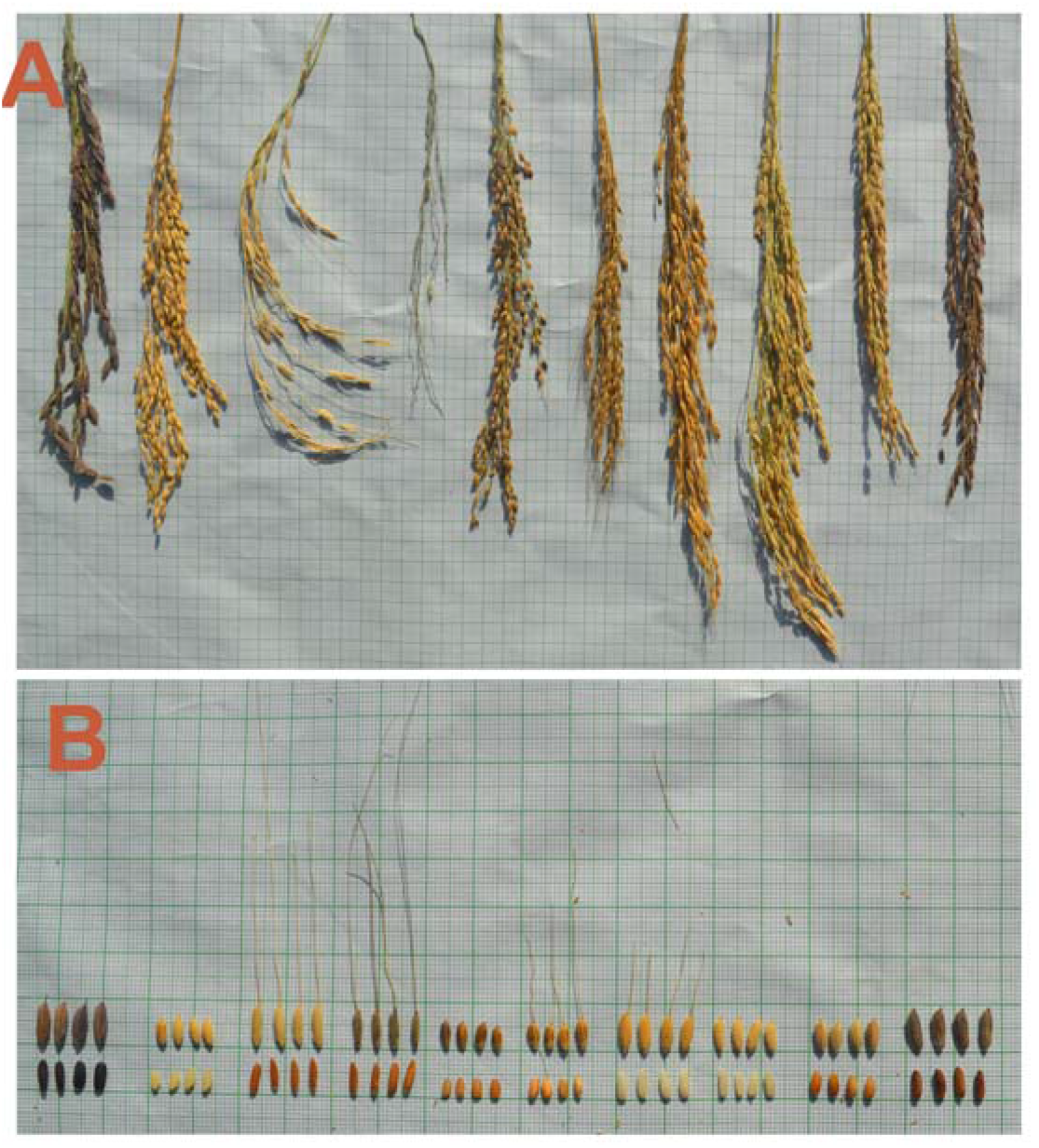
Black rice grain developed through Interspecies hybridization (Pre-breeding). Breeding lines with different grain colour- white, red, brown, black, greenish. Breeding lines (Badshabhog x *O. rufipogon*) showing variability in panicle morphology (Panel-A) and grain shape, size, and grain colour (Panel-B) at F5 population level. From left (Panel A & B)-Local black rice as control for grain colour, parental line *O. sativa* cv. Badshabhog, breeding line BWL1 (semi shattered with awn and red grain), wild rice *O. rufipogon* as donor parental line, then all six breeding lines(non shattered) BWL2, BWL3, BWL4, BWL5, BWL6 and BWL7 (Black).

**Figure 4.**
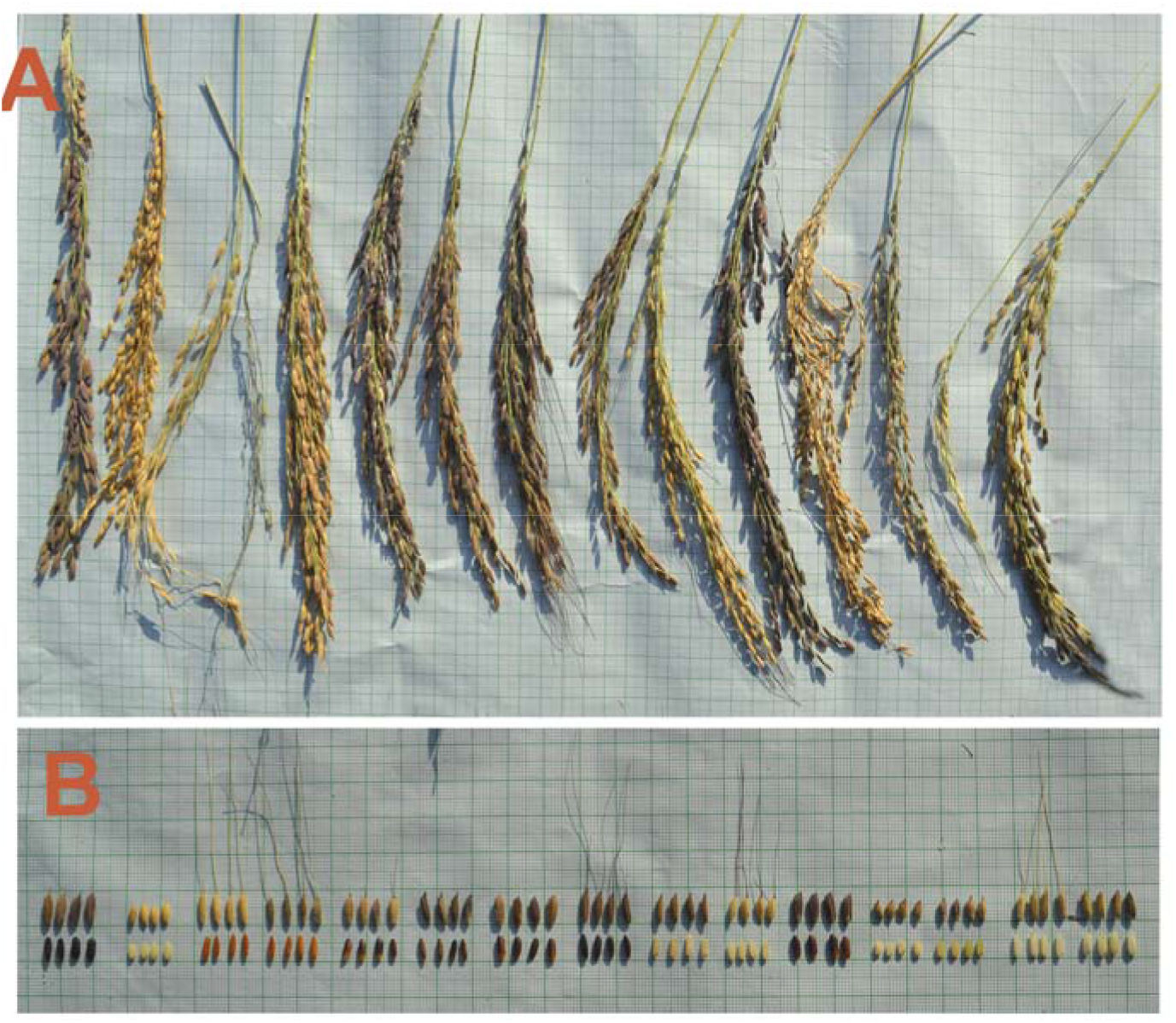
Black rice grain developed through Interspecies hybridization (Pre-breeding). Breeding lines with different grain colour- white, red, brown, black, greenish. Breeding lines (Badshabhog x *O. rufipogon*) showing variability in panicle morphology (Panel-A) and grain shape, size, and grain colour (Panel-B) at F5 population level. From left (Panel A & B)-Local black rice as control for grain colour, parental line *O. sativa* cv. Badshabhog, breeding line BWL8 (red grain with awn and semi shattered), wild rice *O. rufipogon* as donor parental line fully shattered with red grain, breeding lines BWL9-BWL11 (Black grain), BWL12 (Black grain with long awn), BWL13 (white grain without awn), BWL14 (white grain with long awn), BWL15 (black grain without awn), BWL16 (white grain without awn), BWL17 (Greenish grain without awn), and BWL18 (white grain with long awn), BWL19 (white grain without awn).

**Figure 5.**
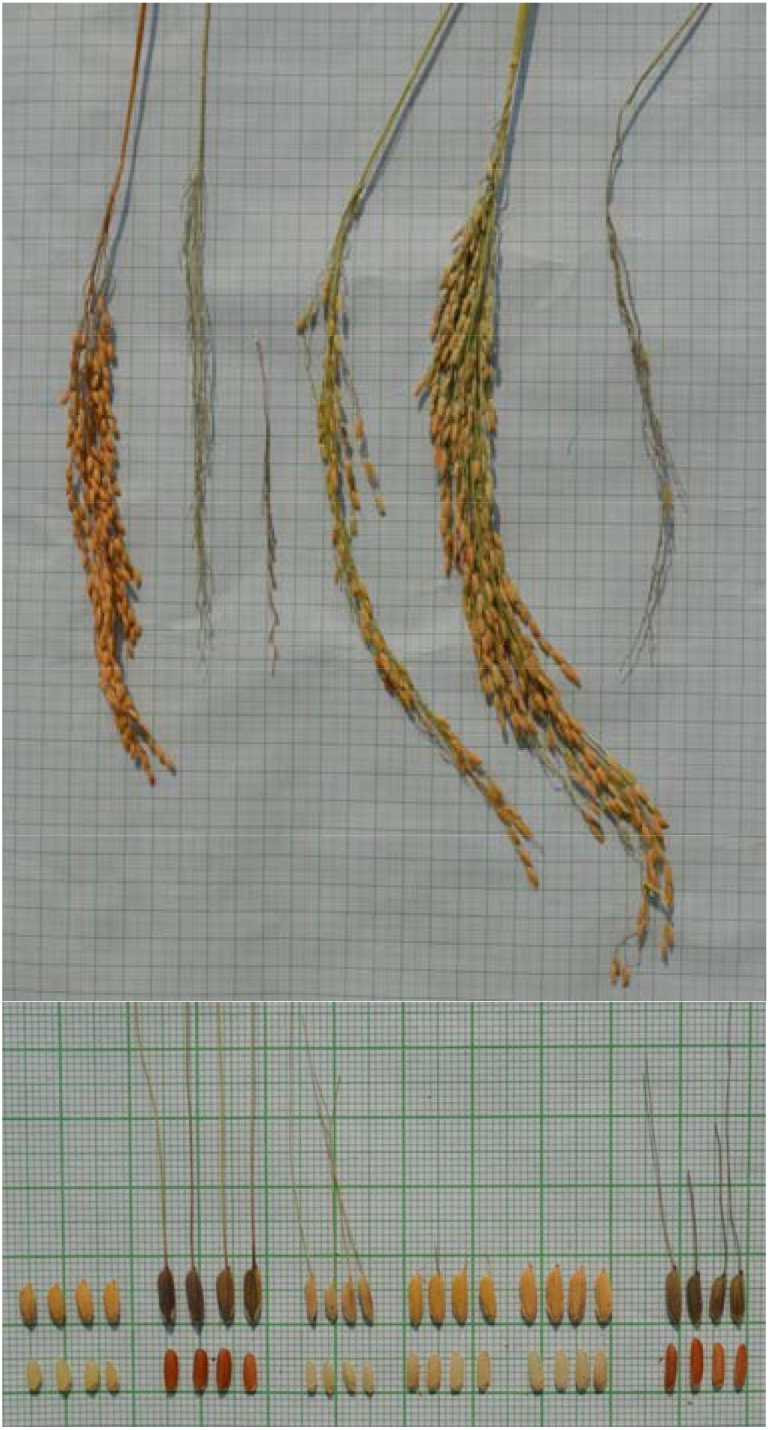
Pre-breeding lines of wide cross (Badshabhog x *O. rufipogon*) showing variability in panicle morphology (Panel-A) and grain shape, size, and grain colour (Panel-B) at F5 population level. From left (Panel A & B)-Parental line *O. sativa* cv. Badshabhog, breeding line BWL20 (red grain with fully shattered long awn), BWL21 (fully shattered and shortest plant height white grain with long awn), BWL22-semi shattered white grain, BWL23 non-shattered white grain, and wild rice *O. rufipogon* as donor parental line red grain.

**Figure 6.**
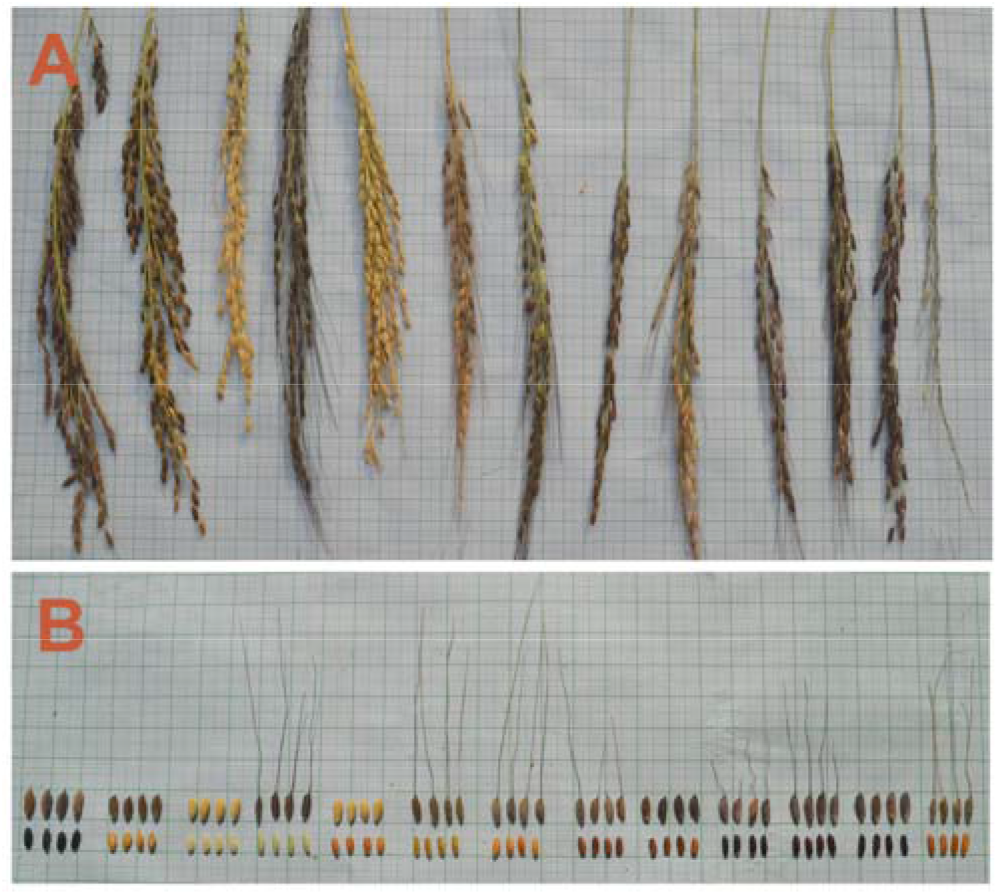
Black rice grain developed through Interspecies hybridization (Pre-breeding). Breeding lines with different grain colour- white, red, brown, black, greenish. Breeding lines (Chenga x *O. rufipogon*) showing variability in panicle morphology (Panel-A) and grain shape, size, and grain colour (Panel-B) at F4 population level. From left (Panel A & B)-Local black rice as control, Chenga-Parent 1 with red grain, breeding lines-CW1-(white grain), CW2-(greenish grain), CW3-(red grain), CW4-(brown grain), CW5-(red grain), CW6-CW7 (Blackish grain), CW8-CW10 (Black grain), and Wild rice *O. rufipogon* (Red grain).

## Discussion

Wide hybridization was performed to broaden the genetic base of the released rice varieties. Released varieties show yield stagnation or saturation due to the narrow genetic base. Breeders always use a few number of parental line to develop the high yielding rice, as a result genetic base is narrowed down. Consequently yield potentiality has not been improved. In the other hand, wild rice is the reservoir of untapped genetic diversity and source of many genes for biotic and abiotic resistance and tolerance. Breeders may utilize these untapped genetic resources for rice improvement through pre-breeding.

In the present study, three wide cross was made between (Badshabhog x *O. rufipogon*), (Chenga x *O. rufipogon*) and (Ranjit x *O. rufipogon*). Breeding lines were analysed according to the DUS test protocol for evaluation of agro-morphological traits and summarized (Table 1–3). Progeny populations of two interspecies hybridization (wide cross) showed black rice grain (Badshabhog x *O. rufipogon*), (Chenga x *O. rufipogon*) at the F_2_ segregating populations but no such black colour grain was observed in the third cross between (Ranjit x *O. rufipogon*). Justification may be like that not all genotype is compatible genetically to induce functional acquired mutation at the genomic level (chromosomal level). Ranjit is a HYV, where many parental genomic combinations have already included in its genome and nonresponsive to structural change at the gene level during meiotic recombination. Other two varieties (Badshabhog and Chenga) are local landrace and conserved by the farmers for many years. Badshabhog is an aromatic but Chenga is a non-aromatic rice cultivar of West Bengal. Frequency of individual plant with black grain was not so high, only few plants with black grain was observed in the F2 populations (Table 1–3, Fig. 3–7). Population size was enough to find out some plants with black grain from the experimental trial field (more than 8000 plant for each cross). Explanation is that the functional acquired mutation mechanism is so rare event that it is not occurring at all the genetic recombination happened during meiotic chromosome pairing and segregation. It also depends on the genome structure of the recipient parent and its maternal components.

**Figure 7.**
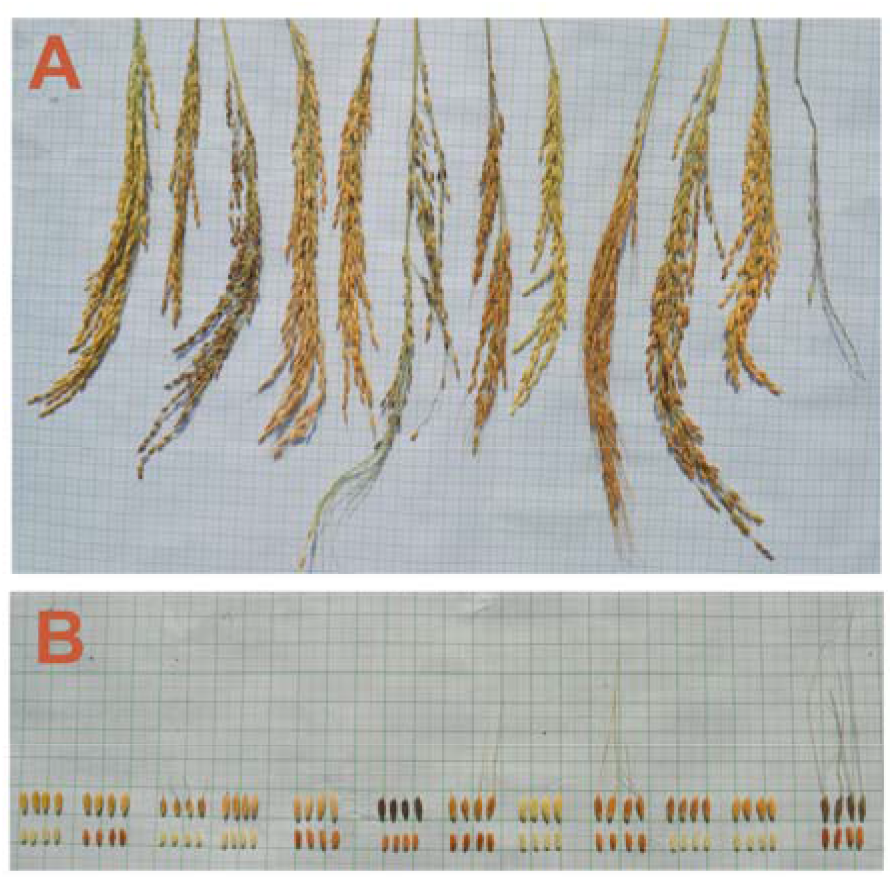
Pre-breeding lines of wide cross (Ranjit x *O. rufipogon*) showing variability in panicle morphology (Panel-A) and grain shape, size, and grain colour (Panel-B) at F5 population level. From left (Panel A & B)- Parental line *O. sativa* cv. Ranjit, breeding line RRL1-(red grain), RRL2-RRL3 (White grain), RRL4-RRL6 (red grain), RRL7 (white grain), RRL8 (Red grain), RRL9-RRL10 (white grain), and wild rice *O. rufipogon* as donor parental line red grain.

Our results of interspecific crossing support the origin of indica type black rice independently taking rearrangement and insertional mutation from wild rice *O. rufipogon* Griff. We propose that indica type black rice of North-East India originated independently through the introgression of necessary genetic components from wild rice and then natural cross breeding in the course of evolution and domestication.

It was predicted that black trait acquired by the rice during or after domestication (Oikawa et al., 2015) based on the study of genome wide genotyping of 21 black rice including other subspecies of rice varieties. Black rice gene can be incorporated still present time at these present eco-climatic conditions/situations. Structural changes in the Kala4 gene, induced ectopic expression of bHLH gene and stimulate the anthocyanin biosynthesis pathway in the pericarp (Oikawa et al., 2015). Results of genome wide genotyping of 21 black rice including red and white rice varieties demonstrated that black rice arose in tropical japonica (Oikawa et al., 2015) and then spread to the indica type subspecies. Evidence support that a small genomic segment of tropical japonica origin found in some indica varieties but not all (Oikawa et al., 2015), containing this Kala4 gene locus. Our present findings not fully agree with this model of black rice origin, that all the indica black rice varieties developed from tropical japonica black rice variety.

In the present study, we report the evidence regarding the black rice development through interspecific hybridization (*O. sativa* cv. Badshabhoh x *O. rufipogon* and *O. sativa* cv. Chenga x *O. rufipogon*). This out crossing initiates the structural change in the Kala4 gene locus specifically at the promoter region which leads to the expression of respective transcription factors gene bHLH. Simultaneously bHLH protein activates the genes associated to anthocyanin biosynthesis in pericarp and developed black grain rice. The newly acquired trait of black rice pericarp was inherited generation after generation in the progeny lines (Table 1–3 and Fig. 3–7). Here, we observed black rice grain in the F_2_ segregating populations and transferring to F3-F4-F5 progeny lines without any genetical defects in the agromorphological traits of developed black rice. Newly developed black rice varieties can be precious genetic material to confirm this insertional mutation in the chromosome 4 during the course of rice evolution. In the present wide breeding investigation, we provide genetic evidence that Kala4 gene, must be changed structurally at the promoter region through the insertion of large genomic segment and induce the expression of bHLH transcription factor gene, which led to the activation of biosynthesis pathway to anthocyanin accumulation to develop black rice pericarp. We support the previous model of the origin and rearrangement of black rice gene Kala4 in tropical japonica (Oikawa et al., 2015) but do not agree that tropical japonica is the only sources of Kala4 gene evolution and black rice development. Here, we propose new model of black rice origin on the basis of the inheritance pattern of newly acquired trait black pericarp in the breeding lines (F2 and onwards) which was not present in any one of the parental lines (Fig. 3–7). Based on the inheritance pattern of the newly acquired black pericarp trait in the breeding populations, we conclude that immediate ancestral progenitor of cultivated rice, *O. rufipogon* is the primary source of black rice pericarp gene and associated regulatory factors, and these were introgressed into the rice varieties through natural out crossing and selection during the course of rice domestication. Introgression of this black pericarp trait either in japonica or indica, shoud be verified through further breeding research.

Previous study showed that black rice landraces exist in at least three subspecies of rice such as indica, tropical japonica and temperate japonica (Oikawa et al., 2015). Model of the black rice birth was proposed based on the genome wide genotyping of 21 black rice varieties including red and white rice varieties, that structural changes in Kala4 gene has been initiated by the insertion of LINE1 (transposon element) near intron2 of the Kala4 gene in the background of tropical japonica rice and considered as single ancestral line of black rice origin. Although, same insertion was found in all type of rice, black, red and white rice, suggest that this insertion spread independently into other subspecies of rice without linking the black trait. Also observed the frequency of insertion was maximum in tropical japonica and a few in indica and aus rice, supporting the model of black rice origin that Kala4 originated from an tropical japonica rice. Subsequently, transferred to the other subspecies of indica and aus. Insertion of LINE1 and partial duplication of LINE1 element was not sufficient to create black rice grain. It was possible to create black rice functional gene Kala4, by insertion of 11.0 kb genome segment which is originated from the upstream −83 kb region of the Kala4 gene. Insertion of 11.0 kb genomic segment was decisive to generate functional allele of black Kala4 gene. There was no such insertion (~11.0 kb genome segment) in any of the red and white rice varieties (Oikawa et al., 2015), only found in black rice varieties. Black rice gene Kala4 originated in the tropical japonica, and then spread to the other subspecies of rice through repeated natural out crossing and artificial selection resulted in refined introgression. Kala4 allele is inherited in a semi dominant manner in the breeding lines. Predicted from the result of 21 black rice genome wide analyses that Kala4 was introgressed many times from japonica to indica subspecies (Oikawa et al., 2015). Our present results partially agree with the proposed model of black rice origin from tropical japonica. Present study of pre-breeding analysis provides important findings about the origin of black rice, from the background of cross between *O. sativa* x *O. rufipogon* and inheritance of alleles associated to the biosynthesis of anthocyanin pigments in rice pericarp (Fig. 3–7). Anthocyanin pigment synthesis is controlled by many genes hence trait is polygenic. We propose one additional new model regarding the origin of black rice. Our experimental breeding analyses support the view that black rice originated from wild rice *O. rufipogon* and introgressed into a refined way into the indica subspecies cultivar Badshabhog and Chenga. Breeding populations showed various type of grain colours–white, red and black, brown and sometimes greenish (Fig. 3–7). Black rice trait is still inherited and maintaining in the F4 and F5 populations without any defective agro-morphological traits at the field trial level (Fig. 2, and Fig. 3–7). Newly developed black rice line represents a valuable source of genetic variation and provides materials to elucidate the genetic basis of black grain pigmentation. Some of the black rice varieties were conserved and grown by the ancient people as a part of their cultural, religious heritage and conserved the most precious heirloom rice landraces.

The evolutionary history of rice anthocyanin biosynthesis genes revealed that the purple leaf trait was negatively selected, and the black rice phenotype is a functional acquired mutation (Oikawa et al. 2015; Zheng et al. 2019). Our present findings are consistent with the previous hypothesis that black rice trait is developed in the rice varieties through functional acquired mutation during the process of cultivation and selection by the ancient people (Oikawa et al., 2015; Zheng et al., 2019). Our present prebreeding results support the view of acquired insertional mutation during the formation of rice varieties with black grain (also called purple). Earlier report was based on genetic and genomic analysis but our present study provides the experimental evidence of construction of new genetic combination at the Kala4 gene locus by inserting large genomic segment at the promoter region to convert white rice to black rice. Rice lines with black grain were observed in the wide crosses between cultivar Badshabhog x *O. rufipogon* and *Chenga x O. rufipogon*. There are no black grain rice in the cross between cultivar Ranjit and *O. rufipogon*. This difference of black pericarp formation indicate that black gene locus Kala4 and other necessary genetic regulator’s construction was depend on genetic architecture of the recipient rice cultivars (*O. sativa*). Ranjit is a high yielding variety released in 1980s but Badshabhog and Chenga are the local rice cultivars available in West Bengal, India. Both the local rice cultivars (Badshabhog and Chenga) have shown more genetic compatibility to reconstruct genetic blue print through rearrangement of chromosomal region (mainly insertion at the promoter region of Kala4 locus) for the biosynthesis of anthocyanin pigment in rice pericarp. This resulted in black (purple) pericarp pigmentation. The results supporting the observation that maternal factors are also responsible for the expression of anthocyanin production in rice grain pericarp. Otherwise the cross between Ranjit x *O. rufipogon*, can also be able to develop black rice pericarp. But progenies of this cross did not show any black grain. Our experimental pre-breeding lines (with black grain) are precious genetic material to the geneticists and rice breeders for the confirmation of earlier views regarding the acquired mutation through insertion of a genetic segment near Kala4 locus. Based on the present experimental results, we propose new hypothesis that black grain colour gene (anthocyanin biosysthesis gene and associated regulatory gene) are inherited from Indian wild rice *O. rufipogon*, and then through natural crossing and artificial selection transferred to indian cultivars (subspecies indica) and independently developed black rice varieties and adopted and evolved in the North-East region of Indian subcontinent. Ultimately, black rice is selected by the ancient people of this region for their nutritional and medicinal value. North-East region of India showed highest genetic diversity among the rice germplasm collected from India (Red rice, Brown rice, black rice and white rice). Therefore, North-East India must be considered as a centre of origin of rice (*O. sativa)*, and indica type rice evolved in this region from the ancestral progenitor wild rice *O. rufipogon* Griff, through domestication and artificial selection. Our present findings also not supporting the earlier proposed hypothesis (Oikawa et al., 2015) based on genetic analysis about the origin of indica type black rice from black rice of tropical japonica type. Because gene flow is constrained by the presence of a complex sterility barriers between the two independently domesticated indica and japonica subspecies of rice (Sweeney et al., 2007). Present results showed that black rice progenies developed through wide crossing between indica type Badshabhog x *O. rufipogon* and Chenga x *O. rufipogon*. Genetic constructions of black rice pigment biosynthesis pathways have been rearranged and reshuffled at the chromosomal level through intergenic recombination in addition to other type of genetic recombinational mutation including intragennic recombination, and all are created in the genetic background of indica type rice varieties (Badshabhog, and Chenga). Our present observations provide a new model that indica type black rice of India originated from such natural wide crossing and artificial selection during the course of evolution of rice, not from black tropical japonica. In spite of several proposed genetical mechanism of black rice formation have been demonstrated by many authors (Oikawa et al., 2015; Sun et al., 2018; Lachagari et al., 2019) there is still a possibility that some additional genes and allelic variants thereof remain to be uncovered (Mbanjo et al., 2020). Based on the review report of earlier author (Mbanjo et al., 2020), here, we believe that black rice trait is newly acquired functional mutation that has been occurred in the present breeding lines, but which had to be happened during the course of rice domestication and evolution. Thus, this is new report about the birth of black rice. Domestication associated genetic mechanisms such as gain-of-function and loss-of-function was involved for the formation of black grain and red grain respectively in the history of rice evolution (Sweeney et al., 2006, 2007). Domestication is not a single event but rather a dynamic evolutionary process that occurs over time (Oikawa et al., 2015; Sun et al., 2018). Evolution of rice domestication is still in debatable conditions (Mbanjo et al., 2020). Therefore, it is possible in the present time suddenly to acquire neofunctionalization genetic mutation near the promoter of Kala4 allele that simultaneously induce the ectopic expression of Kala4 allele to give birth of black rice in the breeding lines of (Badshabhog x *O. rufipogon*) and (Chenga x *O. rufipogon*).

## Conclusion

Our study not only elucidating the origin of black rice through interspecific hybridization but also provides new insights into the rice domestication including the genome evolution in the rice crop. We report the development of aromatic black rice in the history of rice breeding through interspecific hybridization (also called pre-breeding or germplasm enhancenment). Our present evidence supports the model of black rice origin through functional acquired mutation. Genetically it is a rearrangement in the genomic level through chromosomal recombination and insertion of LINE1 element near the Kala4 allele which makes the region instable for partial duplication of LINE1 and additional insertion of 11 kb genomic segment originated from −83kb upstream region of Kala4 gene near the promoter region, which induce ectopic expression of the Kala4 allele and give the birth of Black rice during the history of rice domestication. Present study provides an important genetic basis about the inheritance of black rice gene in the pre-breeding lines that will enable rice breeders and geneticists to effectively exploit wild rice genetic resources for rice improvement. Black rice development through wide hybridization between non-black rice genotypes (*O. sativa* x *O. rufipogon*) certainly helps the understanding of the genetic basis of anthocyanin biosynthesis for the formation of black rice. Here, we propose one additional new model of black rice origin in the genetic background of indica type rice varieties and wild rice (*O. rufipogon*) of india (North-Eastern part of India) as a source of black rice gene. We may predict from this experimental evidence that cultivated rice varieties evolved and domesticated in the North East India where most diversity of rice genotypes prevailed in addition to other region as centre of rice origin. Previous report suggested that black rice originated from tropical japonica during the course of rice evolution and domestication. Rice varieties were developed through natural out crossing and artificial selection involving reined introgression. Most of the varieties are selected by the ancient people/ farmers during the time of domestication and different rice varieties were conserved as their cultural heritage of these heirloom varieties with high nutritional value.

## Acknowledgements

This study was supported by the University of North Bengal. SCR is thankful to the university authority for giving me this financial support to do the inter-specific hybridization in the Plant Genetics & Molecular Breeding Laboratory, Department of Botany, University of North Bengal, India.

## Contributions

**SCR**-conceptualized and carried out the interspecific hybridization to broaden the genetic base of the cultivated varieties, identified wild rice genotype *O. rufipogon* from Raiganj, West Bengal and used in this out crossing breeding, planned and designed the breeding material for present study, wrote and reviewed the whole manuscript, arranged all the figures in the manuscript. **PS**-provided tables with data analysis and carried out breeding analysis.

